# The effect of developmental pleiotropy on the evolution of insect immune genes

**DOI:** 10.1101/2021.05.12.443901

**Authors:** Thi Minh Ngo, Alissa M. Williams, Ann T. Tate

## Abstract

The pressure to survive relentless pathogen exposure explains the frequent observation that immune genes are among the fastest-evolving in the genomes of many taxa, but an intriguing proportion of immune genes also appear to be under purifying selection. Though variance in evolutionary signatures of immune genes is often attributed to differences in gene-specific interactions with microbes, this explanation neglects the possibility that immune genes participate in other biological processes that could pleiotropically constrain adaptive selection. In this study, we analyzed available transcriptomic and genomic data from *Drosophila melanogaster* and related species to test the hypothesis that there is substantial pleiotropic overlap in the developmental and immunological functions of genes involved in immune signaling and that pleiotropy would be associated with stronger signatures of evolutionary constraint. Our results suggest that pleiotropic immune genes do evolve more slowly than those having no known developmental functions, and that signatures of constraint are particularly strong for pleiotropic immune genes that are broadly expressed across life stages. However, pleiotropic immune genes also contain a significantly higher proportion of positively selected sites and substitutions are more likely to be under positive selection, suggesting a mechanism to circumvent evolutionary constraint. These results support the general yet untested hypothesis that pleiotropy can constrain immune system evolution, raising new fundamental questions about the benefits of maintaining pleiotropy in systems that need to rapidly adapt to changing pathogen pressures.

## Introduction

Over evolutionary time, organisms have developed defense mechanisms against microbial pathogens and parasites which counter-adapt, in turn, to maintain successful infection strategies. Host immune systems put selective pressure on microbes to evade host recognition, repel antimicrobial effectors, and even manipulate immune signaling components to dampen host defenses (Schmid-Hempel 2008; Heil 2016). Hosts that cannot circumvent these mechanisms could suffer massive fitness costs from infection. As a result, pressure from pathogens and parasites represents a major driving force in molecular evolution (Paterson, et al. 2010).

How should we expect selection to act on immune system genes? Host adaptation to microbial pressure should drive positive, directional selection or, in the face of coevolutionary negative frequency dependence, balancing selection that maintains polymorphism in populations (Casals, et al. 2011; Sackton 2019). Studies in species as diverse as humans (Mukherjee et al. 2009; Casals et al. 2011), non-human mammals (Seabury, et al. 2010; Areal, et al. 2011) and insects (Sackton, et al. 2007; Obbard, et al. 2009; Rottschaefer, et al. 2015) have found evidence for both positive and balancing selection in immune system recognition and effector genes (Unckless, et al. 2016). For example, Obbard *et al*. found that *Drosophila melanogaster* immune genes, as a class, have higher rates of adaptive substitution than location-matched non-immune genes (Obbard, et al. 2009). However, these trends were driven by a few particularly rapidly evolving genes associated with a subset of immune signaling pathways, while purifying selection was surprisingly prevalent on immune genes in other pathways. If parasites frequently target or evade signaling components, why wouldn’t those targets show rapid adaptation?

The answer may depend on a crucial but underappreciated quality of immune systems. Genetic pleiotropy arises when a single gene product contributes to multiple discrete phenotypic traits, and many components of immune pathways appear to be pleiotropic. Since the discovery of the Toll pathway, for example, numerous studies (and indeed Nobel prizes) have recognized its conserved dual role in development and innate immune system signaling (Lemaitre, et al. 1997; DiAngelo, et al. 2009; Anthoney, et al. 2018), and proposed that this could impose constraints on immune system evolution (Obbard, et al. 2009; Tan, et al. 2021). More broadly, a recent study estimated that ∼17% of human genes affect multiple discrete phenotypic traits, and functional enrichment analysis of this pleiotropic gene set revealed immune system functions to be among the most over-represented processes (Sivakumaran, et al. 2011). When a pleiotropic mutation affects uncorrelated traits, opposing forces of selection on each trait can reduce the efficacy of selection and resist the fixation of adaptive substitutions (Fraïsse, et al. 2018). Thus, the adaptive evolution of pleiotropic immune genes may be constrained by the deleterious effects of substitutions on other traits.

Pleiotropy between development and immunity is particularly intriguing because a developmental program must be carried out faithfully for an organism to progress through its life cycle, resulting in purifying selection on genes involved in embryonic and early life development. Indeed, developmental pleiotropy (defined by the number of genetic interactions (Stark, et al. 2006)) has been shown in *D. melanogaster* to constrain positive selection in early-expressed genes due to a higher number of functional interactions in those genes that render mutations deleterious (Artieri, et al. 2009). We hypothesize that developmental pleiotropy could constrain immune gene evolution, particularly for genes involved in the most complex stages of development (Tian, et al. 2013), leading to an under-representation of signatures of positive selection on immune genes relative to theoretical expectations.

Insects can serve as particularly valuable models for studying the evolutionary consequences of developmental and immunological pleiotropy due to their discrete life stages, a wealth of genomic resources, and availability of studies on immune gene function (Consortium 2013; Palmer and Jiggins 2015; Viljakainen 2015). The canonical components of an insect innate immune response include microbial recognition, signal transduction to initiate cellular and humoral responses, and production of effector molecules for pathogen clearance (Lemaitre and Hoffmann 2007). Many genes and signaling pathways previously identified as core participants in these processes are also broadly conserved among species (Waterhouse, et al. 2007), including two of the best studied pathways, Toll and Imd, which coordinate expression of antimicrobial peptides and other pathogen-clearing effectors (Ferrandon, et al. 2007; Tanji, et al. 2007). While the Toll pathway is the most recognized example of developmental and immunological pleiotropy in insect immune systems, previous work has highlighted potential pleiotropy within other pathways (Tate and Graham 2015). For example, the same components of the melanization pathway responsible for tanning the insect cuticle after each larval molt are also used for melanizing parasitoid eggs and neutralizing pathogenic fungi, leading to allocation issues when an insect needs to accomplish both at once (McNeil, et al. 2010; Parker, et al. 2017). Thus, pleiotropy is likely to interfere with the deployment of immune responses if a host needs to use a gene product for both development and immunity in the same life stage. Even if these functions are segregated into different life stages, however, could pleiotropy still constrain immune system evolution?

We predict that immune genes that have a pleiotropic developmental function will be more likely to experience evolutionary constraint, as defined by slower rates of evolution and a lower frequency of positive selection, than immune genes that have no known developmental function. Further, we predict that pleiotropic genes that are crucial to multiple developmental stages will be the most constrained, relative to genes involved in more specific and less conserved developmental processes. To investigate these predictions, we combine transcriptional and functional genomics data from fruit flies (*Drosophila* spp.) to characterize the overall and immune pathway-specific degree of pleiotropy among immune and developmental genes. We then analyze the rates of evolution in immune genes using genomics data from 12 sequenced *Drosophila* species. Empirical support for our predictions would raise the question of why evolution would maintain pleiotropy between development and immunity given the potential for conflict and constraint. On the other hand, if pleiotropic immune genes are not more constrained than non-pleiotropic ones, this study could inspire future investigations into compensatory evolution and the role of network architecture in minimizing evolutionary conflict.

## Results

### Extent of developmental pleiotropy in immune genes

To determine the prevalence of developmental pleiotropy among immune genes, we started by curating separate lists of immune and developmental genes. Previous studies have employed various methods to curate gene lists, ranging from using only Gene Ontology annotations (Fraïsse, et al. 2018) to compiling experimentally confirmed and/or computationally predicted immune gene orthologs (Early, et al. 2017). Taking these different approaches into account, we employed several sources to assemble a comprehensive suite of genes that participate in immunity (Table 1 and Methods). In total, we assembled a list of 808 immune genes, of which 551 genes have known canonical roles in immunity and 107 genes play a role in immune system development, as annotated by Gene Ontology and previous studies (Early, et al. 2017). The degree of overlap between different immune gene list sources can be found in Supplemental Figure 1. The list of developmental genes contains 3346 genes, of which 262 genes are annotated specifically as “embryonic development” genes and 508 as “post-embryonic development.” Some embryonic development genes also participate in post-embryonic development (overlap visualized in Supplemental Figure 2).

**Table 1:**
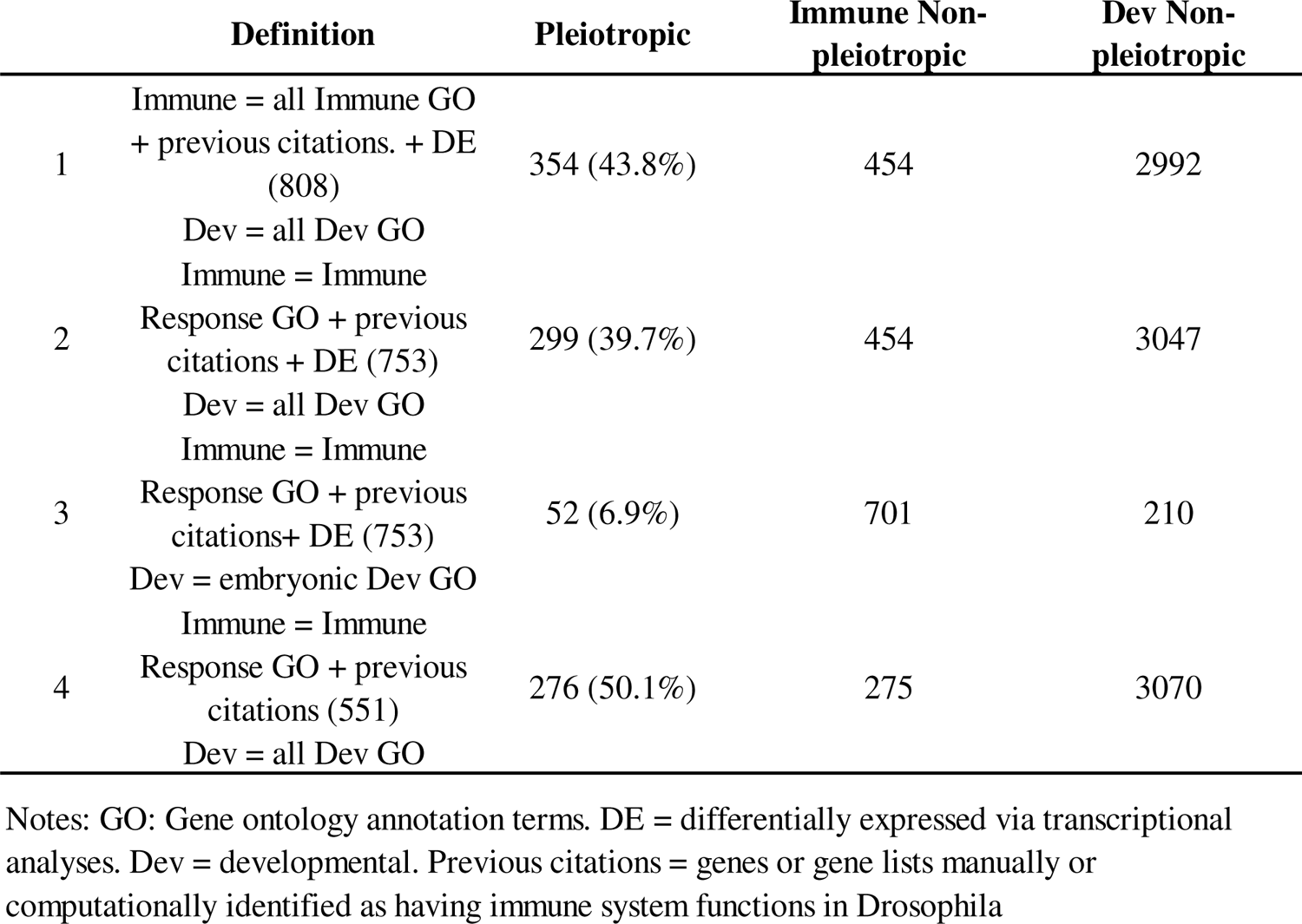
The extent of pleiotropy as defined with different annotation methods.

Genes that appear in both the immune and developmental gene lists were labeled as “pleiotropic.” When considering immune genes as those identified by all methods including manually curated, GO annotated and differentially expressed genes, we found 354 immune genes (43.8%) to be pleiotropic (Table 1, row 1). When constraining the definition of immune gene to those that directly contribute to an immune response while excluding genes participating in development of the immune system, 299 (39.7%) genes are considered pleiotropic (Table 1, row 2). Under the most conservative definition of development (only genes that directly participate in embryonic development or 7.8% (262/3346) of all annotated developmental genes), 52 immune genes (6.9%) still meet the definition of pleiotropy (Table 1, row 3). The full list of immune, developmental, and pleiotropic genes under different categorization methods is included in Supplemental Table 1. Note that although we used several methods to compile a list of pleiotropic genes, the conclusions generated throughout this study are robust to different categorical definitions of immunity, development, and pleiotropy (see expanded discussion in Methods). Therefore, from this point on, for simplicity, we refer to our immune gene group as those defined using the sources from Table 1, row 2, which comprises Immune Response GO-annotated genes, immune genes employed in previous large scale studies, and a core set of genes differentially expressed in ten bacterial infections (Troha et al. 2018).

### Comparison of pleiotropic and non-pleiotropic immune gene characteristics

Immune genes can be categorized into different classes, such as recognition, signaling, and effector, depending on their canonical function in an immune response. We were curious whether certain classes of immune genes are more likely to have pleiotropic status than others. We divided immune genes into major categories, relying on both annotation from previous studies (Sackton, et al. 2007; Early, et al. 2017) and manual annotation based on gene description in FlyBase (Supplemental Table 2). According to this classification system, the number of genes confirmed to each category includes 33 recognition genes, 123 signaling genes and 27 effector genes (Supplemental Figure 3). As represented in Figure 1A, the signaling immune class contains the highest proportion of pleiotropic genes (66.67%, n = 123), and the different groups contain a significantly different proportion of pleiotropic genes overall (X^2^= 37.94, p < 0.0001). Moreover, using the PANTHER pathway database, we found that pleiotropic genes are, on average, associated with more pathways than non-pleiotropic ones (Supplemental Table 3).

**Figure 1.**
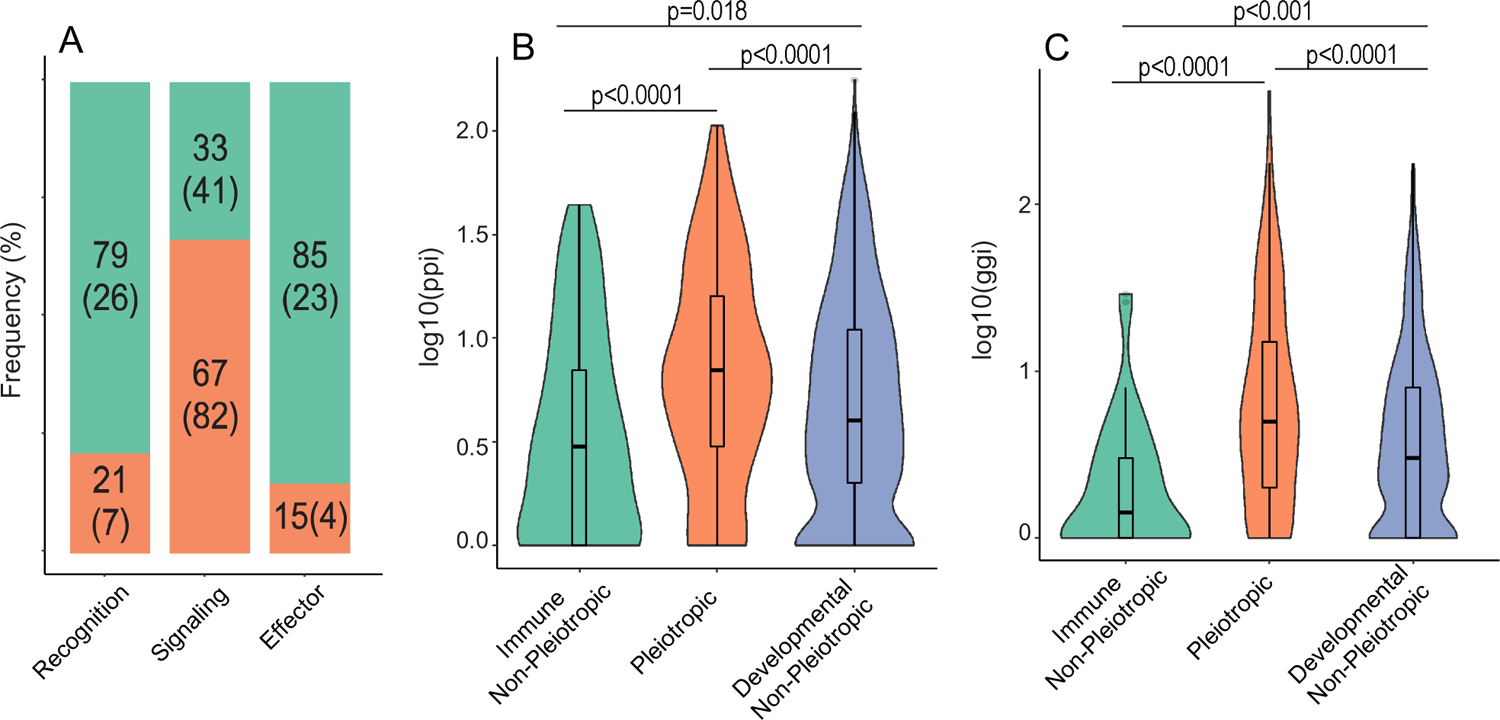
Overall characterization of pleiotropic and non-pleiotropic immune genes. Each immune gene was assigned a “gene class” (**A**) depending on their canonical function in an immune response. For each class, the percentage of pleiotropic (those with developmental roles; pink) and non-pleiotropic genes (green) was determined (big number: proportion; number in parentheses: number of genes in that category). The number of known protein-protein interactions (ppi; **B**) and number of known gene-gene interactions (ggi; **C**) were also calculated for genes annotated as immune non-pleiotropic (green), pleiotropic for development and immunity (pink), or developmental non-pleiotropic (blue), represented on a log-scale and statistically analyzed using Kruskal-Wallis tests for overall significance followed by post-hoc pairwise Dunn tests (Benjamini-Hochberg-adjusted p values on figure).

We also wanted to know whether our curated immune-developmental pleiotropic genes exhibit characteristics associated with alternative definitions of pleiotropy, such as a high number of associated protein-protein interactions and gene-gene interactions that reflect activity at the molecular level. When comparing pleiotropic and non-pleiotropic immune genes (Figure 1B-C), we do find that pleiotropic genes have significantly more protein-protein interactions (Kruskal-Wallis w/ Dunn post-hoc test, p.adj = 3.8e-05) and more gene-gene interactions (Kruskal-Wallis w/ Dunn post-hoc test, p.adj = 6.3e-07). Moreover, pleiotropic genes are associated with more Biological Processes (Wilcoxon test, p < 2e-16) and Molecular Functions (Wilcoxon test, p < 2e-16) GO terms than non-pleiotropic genes (Supplemental Figure 4).

### Expression specificity across stages and tissues between pleiotropic and non-pleiotropic genes

To investigate the hypothesis that broadly expressed pleiotropic genes are under stronger evolutionary constraint than specific ones, we determined gene expression specificity across life stages and tissues for pleiotropic and non-pleiotropic immune genes using the τ specificity index ((Yanai, et al. 2005), see methods). A large τ value indicates specific expression while a small value indicates broad expression across stages or tissues. While we could not confidently determine whether any given gene plays only a developmental or immunological role or both at any given stage, genes involved in development at multiple life stages may present a temporal as well as evolutionary constraint on the immunological function of that gene.

We found that, in uninfected insects, pleiotropic immune genes (median τ = 0.670) are significantly more broadly expressed across stages than non-pleiotropic immune genes (Figure 2A, median τ = 0.731; Kruskal-Wallis w/Dunn test, p.adj = 0.0009), but have similar expression breadth profiles as non-pleiotropic developmental genes (median τ = 0.691; Kruskal-Wallis w/Dunn test, p.adj = 0.07). We also found that the most stage-specific pleiotropic genes, determined by the top quartile in τ value, disproportionately exhibit maximal expression during the embryonic stage (43% among specific pleiotropic genes vs 3.6% among specific non-pleiotropic immune genes) while the most specific non-pleiotropic immune genes exhibit a relatively even distribution of maximal expression across subsequent stages (Figure 2B, Supplemental Table 4). At the tissue level, pleiotropic genes are also expressed more broadly than non-pleiotropic immune genes, and this trend is consistent throughout all life stages (Figure 2C). We found no significant differences in tissue expression specificity between developmental genes and pleiotropic genes except in the adult stage (Figure 2C), where developmental genes showed more specific patterns of expression.

**Figure 2.**
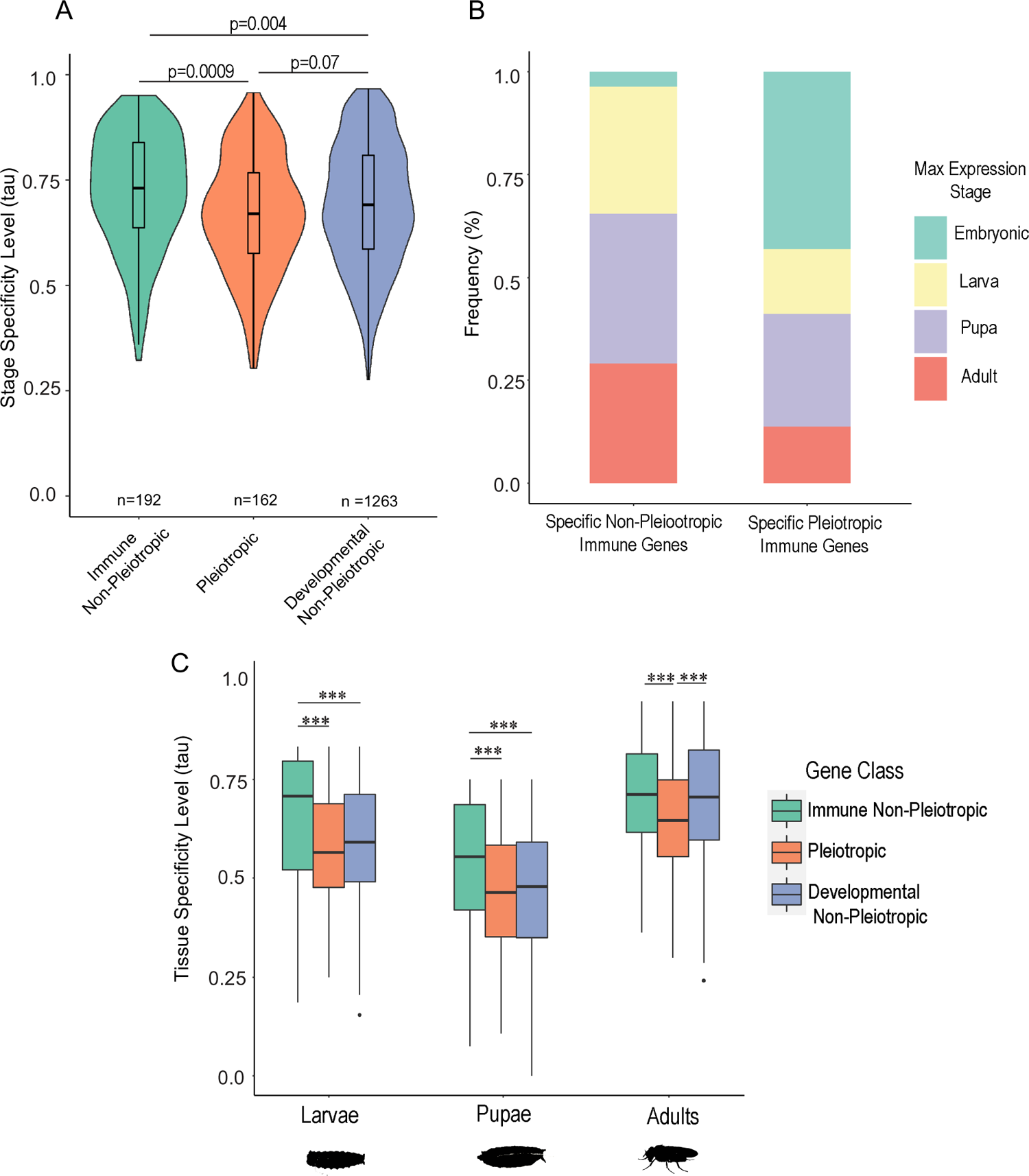
Comparison of relative life stage and tissue specificity of gene expression among immune, developmental, and pleiotropic genes. The stage specificity tau value, which varies from 0 (broadly expressed across all stages) to 1 (expressed in only one stage) was calculated for genes within each class (**A**). For the non-pleiotropic and pleiotropic immune gene group (**B**), the genes within the top 25th percentile of τ value were characterized as “specific genes”, and the stage with the highest expression for each gene was determined and tallied for the whole group. To compare tissue gene expression specificity between pleiotropic and non-pleiotropic genes within each life stage (**C**), the tau value (tissue specificity level) was calculated for each gene across tissues. Differences among groups were statistically analyzed using Kruskal-Wallis tests for overall significance followed by post-hoc pairwise Dunn tests (Benjamini-Hochberg-adjusted p values on figure; *** indicates p.adj < 0.001).

### Evolutionary rates among different gene categories

To address whether pleiotropic genes are more evolutionarily constrained than non-pleiotropic genes, we calculated *d_N_/d_S_* values using codeml site model M0 in PAML v4.9j (Yang 2007), which assigns a single *d_N_/d_S_* value to an entire tree (see Methods). We ran this PAML model for concatenations of genes in 12 *Drosophila* species (Supplemental Table 5), where each concatenation represented one of three categories of genes: non-pleiotropic immune, pleiotropic, and non-pleiotropic developmental. Genes for each concatenation were defined using Table 1 row 2, and after quality control, these concatenations contained 356, 231, and 2067 genes, respectively. We also ran codeml site model M0 on each individual gene included in the concatenations; these model runs were successful for 348 non-pleiotropic immune genes, 227 pleiotropic genes, and 2037 non-pleiotropic developmental genes (see Methods).

The model runs on the concatenated gene lists yielded *d_N_/d_S_* estimates of 0.098 for non-pleiotropic immune genes, 0.077 for pleiotropic genes, and 0.078 for non-pleiotropic developmental genes. Meanwhile, model runs on individual genes yielded median *d_N_/d_S_* estimates (Figure 3A) of 0.085, 0.063, and 0.063 respectively, and these three categories exhibited significantly different *d_N_/d_S_* distributions based on a Kruskal-Wallis test (Figure 3A, χ² = 66.53, p = 3.57e-15). Pairwise comparisons of individual gene *d_N_/d_S_* values were calculated using post-hoc Dunn tests adjusted for multiple comparisons. The comparison between pleiotropic genes and developmental non-pleiotropic genes does not show a statistically significant difference (p = 0.95), but non-pleiotropic immune genes have a statistically different distribution relative to both non-pleiotropic developmental genes (p = 1.8e-15) and pleiotropic genes (p = 4.2e-08).

**Figure 3.**
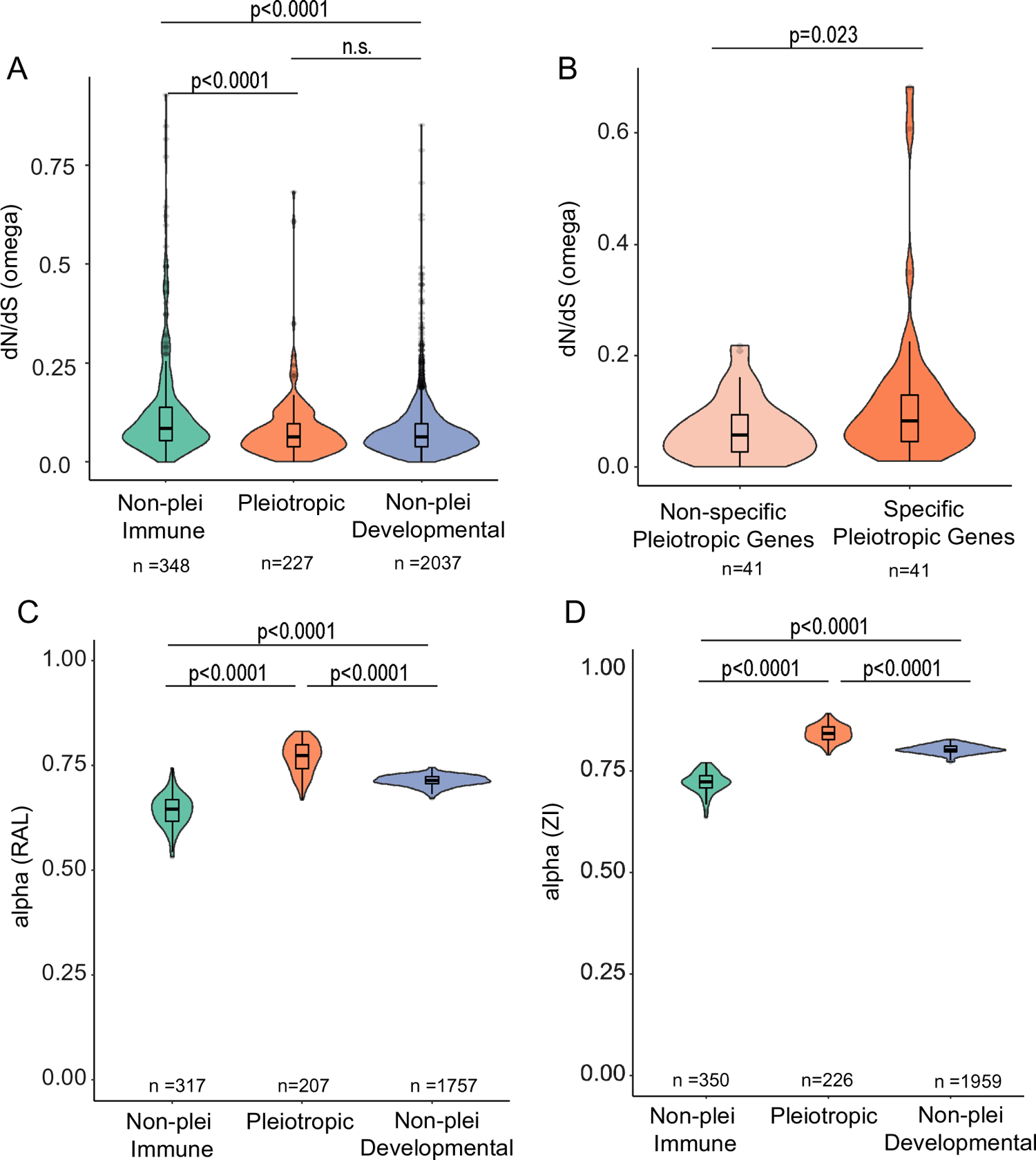
Associations between genetic pleiotropy, stage specificity, and dN/dS ratios. dN/dS values (A) were compared among non-pleiotropic immune genes, genes with pleiotropic roles in development and immunity, and developmental genes with no known pleiotropic role in immunity. dN/dS values were also compared between pleiotropic genes that scored within the top and bottom quartiles of stage-specific expression (B), where non-specific pleiotropic genes are broadly expressed across life stages (tau <=0.576) while the top quartile are specifically or maximally expressed in fewer stages (tau >=0.767). Differences among groups were statistically analyzed using a Kruskal Wallis test (A) followed by post-hoc Dunn tests (p values BH-adjusted) or a Wilcoxon test (B). P values reproduced on the figure; n.s. = not significant (p.adj > 0.05). C and D depict *a* values for all three gene categories in two populations of *Drosophila melanogaster*, Raleigh (RAL) and Zambia (ZI). *a* values were calculated using MultiDFE on 100 bootstrap replicates of summed site frequency spectra (SFS) for each gene category. Distributions were compared using a Kruskal-Wallis test followed by post-hoc Dunn tests in R.

Within the pleiotropic gene set, we found that the most specifically stage-expressed genes (top τ quartile, e.g. Fig. 2B) had significantly lower *d_N_/d_S_* ratios than the most broadly expressed pleiotropic genes (bottom τ quartile; n = 41/quartile, Wilcoxon test, p = 0.023, Fig. 3B).

### Evidence for positive selection across gene categories

To determine whether there is evidence for positive selection in any of the three gene categories, we ran codeml site models M7 and M8 in PAML v4.9j (Yang 2007) on each concatenation (see Methods). Model M7 splits the codons in the alignment into 10 groups, where each group contains 10% of the full alignment and has a *d_N_/d_S_* value constrained to be less than 1. Model M8 splits the alignment into 11 groups, where the proportion of the alignment represented by each group varies; the first 10 groups in M8 have *d_N_/d_S_* values constrained to be less than 1, while group 11 can have a *d_N_/d_S_* value greater than 1 (representing positive selection in that group of codons). These two models are compared using a likelihood ratio test with two degrees of freedom to determine whether a model allowing for positive selection is a better fit for the data than a model that does not.

A likelihood ratio test between the two models provided significant evidence for positive selection in a fraction of sites within the concatenated alignments of each of the three categories (p < 0.001 for all). In the case of the non-pleiotropic immune gene concatenation, the proportion of sites in the eleventh category was 0.007 with an omega value of 5.37. The proportion of sites in the eleventh category for the pleiotropic gene concatenation was 0.015 with an omega value of 1.37. The non-pleiotropic developmental gene concatenation yielded a similar result as the pleiotropic one, with a proportion of 0.018 and omega value of 1.29. The three proportions calculated by model M8 were all statistically different from one another (Chi-squared = 1034.6, p < 2.2e-16) and each pairwise comparison of proportions was statistically different even after Bonferroni correction (p < 2.2e-16 for all three).

We also compared proportions of sites under positive selection identified by Bayes Empirical Bayes (BEB) analysis in model M8. We defined positively selected sites as those with a probability of *d_N_/d_S_* > 1 of 0.95 or above as detected by BEB analysis. The percentages of sites under positive selection were 0.059%, 0.12%, and 0.13% for the non-pleiotropic immune, pleiotropic, and non-pleiotropic developmental categories, respectively. A Chi-squared test showed that these proportions were statistically different from one another (p = 8.99e-12). Pairwise Chi-squared tests with Bonferroni correction confirmed the statistically significant difference between the non-pleiotropic immune concatenation and the other two concatenations (p < 0.001 in both cases). There was not a significant difference in this proportion between the pleiotropic and non-pleiotropic developmental concatenations (p = 0.40).

### Evidence of adaptive evolution across gene categories

The PAML results indicated that non-pleiotropic immune genes had higher *d_N_/d_S_* values than either pleiotropic genes or non-pleiotropic developmental genes; the latter two categories were not statistically different from one another (Figure 3A). To help determine whether this difference in *d_N_/d_S_* values was driven by adaptive evolution and/or relaxed selection, we used MultiDFE to calculate α *and* ω_a_ for 100 bootstrap replicates of each of the three categories separately for two populations of *Drosophila melanogaster*: Raleigh (RAL) and Zambia (ZI). We obtained site frequency spectra from PopFlyData in the iMKT package (Murga-Moreno, et al. 2019) and final values of α and ω*_a_* were determined using a Jukes-Cantor correction.

We found that there were significant differences in α across categories in both the RAL and ZI populations (Figure 3C,D; p < 2.2e-16 for both). Median values of α for the non-pleiotropic immune genes, pleiotropic genes, and non-pleiotropic developmental genes, respectively, were 0.647, 0.774, and 0.714 for RAL and 0.724, 0.843, and 0.803 for ZI. Post-hoc Dunn tests revealed that there were significant differences in all pairwise comparisons of α for both populations even after Bonferroni correction (p < 0.001 in all cases). For both populations, the median α value was highest in the pleiotropic gene class, followed by the non-pleiotropic developmental gene class and then by the non-pleiotropic immune gene class.

There were also significant differences in ω_a_ across categories in both populations (Supplemental Figure 5; p = 3.202e-13 for RAL, p < 2.2e-16 for ZI). Median values of ω_a_ for non-pleiotropic immune genes, pleiotropic genes, and non-pleiotropic developmental genes, respectively, were 0.160, 0.178, and 0.152 for RAL and 0.192, 0.214, and 0.187 for ZI. Post-hoc Dunn tests for the RAL population found that all pairwise comparisons of ω_a_ were significant after Bonferroni correction (p < 0.001 in all cases). For the ZI population, ω_a_ was significantly different between the pleiotropic category and both other gene classes even after Bonferroni correction (p < 0.001 for both) but was not significantly different for the non-pleiotropic immune vs. non-pleiotropic developmental comparison (p = 0.0660).

### Evidence of positive selection in immune signaling pathways

The high overall frequency of pleiotropy among immune signaling genes (Figure 1A) prompted us to examine the distribution of *d_N_/d_S_* along the three major insect immune signaling pathways (Figure 4: Imd, Toll, Jak/STAT) to further investigate whether there are certain components that tend to be pleiotropic or show discernable patterns of ω values. We also ran codeml site models M7 and M8 in PAML on these individual pathway components to determine whether any harbored strong evidence of positive selection.

**Figure 4.**
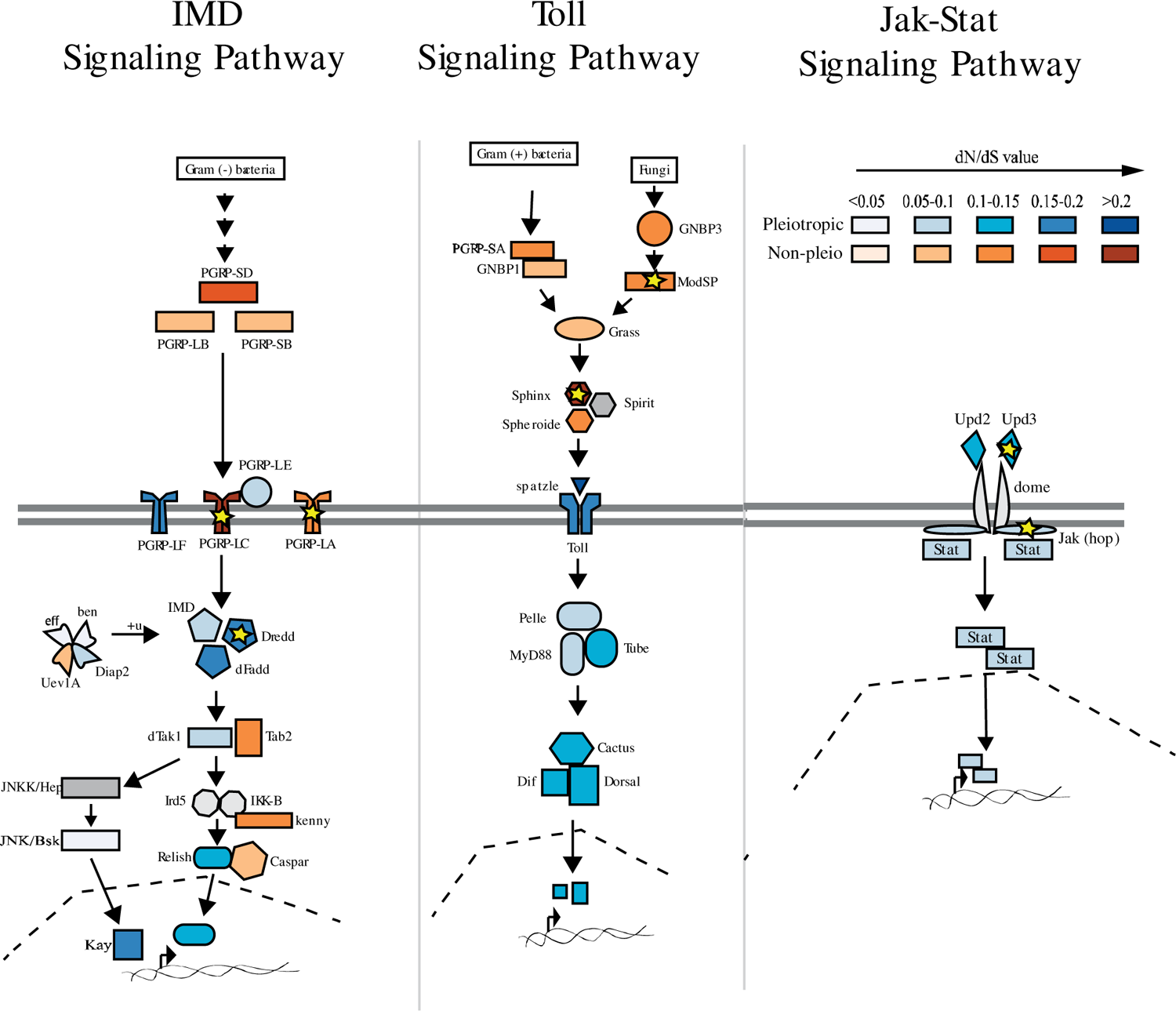
Examining the pleiotropy status and dN/dS levels for genes participating in major insect immune signaling pathways. The color indicates whether has pleiotropic roles in development and immunity (blue) or functions exclusively in immunity (orange). Each color is shaded according to dN/dS level of each gene, with the darker shade represent a higher ω value within the gene’s respective pleiotropic or non-pleiotropic group. Pathway components reflect annotated genes from KEGG. Components for which no pleiotropy status available (e.g. JNKK, Spirit) are shown in gray. Yellow stars indicate genes that have a positively selected fraction of sites (dN/dS > 1) as determined by comparison of PAML models M7 and M8 outputs (see methods).

As illustrated in Figure 4, extracellular signaling components tend to be non-pleiotropic, while intracellular signaling components are consistently pleiotropic. The exceptions are immune-specific adapters within the IMD signaling pathway (e.g. Tab2 and Kenny) that interact with pleiotropic proteins. There were no clear patterns with regard to overall *d_N_/d_S_* distribution along these pathways, as both the intracellular and extracellular compartments contain proteins with relatively low and high *d_N_/d_S_* values. While any estimate of positive selection for individual genes through comparison of model M7 and M8 outputs will be underpowered and thus overly conservative because of the small number of sites, our analysis did still identify several genes in these pathways that contain sites undergoing positive selection (Figure 4 gold stars). Most of these genes are extracellular (ModSP, Sphinx, Upd3) or involved in pathogen recognition (e.g. PGRP-LC; PGRP-LA) and thus conform to the typical profile for immune genes experiencing rapid evolution. However, we also found that the pleiotropic intracellular caspase Dredd exhibited statistical evidence of positive selection (Model 8: 5.7% of sites with average ω= 1.22, p = 0.0006), providing a salient candidate for future studies of pleiotropy.

## Discussion

Researchers have long recognized that some immune genes, such as those in the Toll pathway, play double-duty in development (Lemaitre, et al. 1996), and posited that it might constrain immune system evolution (Obbard, et al. 2009). Pleiotropy seems like it would be a liability for a host, for multiple reasons – what if a gene product cannot be deployed to fight a parasite because it is already being fully allocated to development? Shouldn’t purifying selection on developmental genes constrain the rate of adaptation against parasite pressure, putting the host at a disadvantage during coevolution with rapidly evolving parasites? In this study, we investigated the relationship between immunity-development pleiotropy and signatures of molecular evolution in *D. melanogaster* immune genes. Our results provide clear quantitative evidence for the notion that pleiotropy between development and immunity is actually quite common (Tate and Graham 2015). Moreover, immune genes involved in development exhibit stronger signatures of evolutionary constraint than non-pleiotropic immune genes, particularly if they are broadly expressed across life stages, consistent with our hypothesis of evolutionary constraint.

While overall *d_N_/d_S_* values are lower for pleiotropic immune genes, our analyses looking specifically for evidence of positive selection suggest that twice as many sites are under positive selection in pleiotropic vs. non-pleiotropic immune genes, raising the question of whether compensatory evolution at specific sites might play a role in relieving evolutionary antagonism between development and immunity. Interestingly, pleiotropic genes were not significantly different from non-pleiotropic developmental genes in terms of *d_N_/d_S_* values (Figure 3A). This observation suggests that genes with both immune and developmental functions are similar to developmental-only genes rather than immune-only genes (or an intermediate between the two groups) in terms of evolutionary constraint. We also found that among the three gene categories, pleiotropic immune genes had the highest α values while non-pleiotropic immune genes had the lowest α values (Figure 3C,D), suggesting that increased *d_N_/d_S_* values in the non-pleiotropic immune category are at least partially due to an increase in relaxed selection relative to the pleiotropic category. A higher proportion of adaptive substitutions in the pleiotropic category is consistent with the stronger purifying selection in those genes compared to non-pleiotropic immune genes.

Our systematic curation of transcriptional data, GO terms, and functional evidence from *D. melanogaster* revealed that about 40-44% of immune genes are pleiotropic with development. This estimate aligns with a phenotypic screening study in mammals that more generally classified approximately 65% of screened alleles as pleiotropic across a range of phenotypes (De Angelis, et al. 2015). Upon analyzing the different immune gene classes for their prevalence of pleiotropy (Figure 1A), we found that immune signaling genes are most likely to participate in developmental functions. This is expected since a signaling pathway is capable of activating the transcription of multiple genes, as opposed to, for example, effector genes which likely only interact with microbial pathogens or have specific immune functions. Further, genes annotated as pleiotropic through our classification method also exhibited significantly higher values of molecular parameters associated with pleiotropy (Alvarez-Ponce, et al. 2017), as they have more protein-protein and gene-gene interactions (Figure 1B,C) and are expressed more broadly across life stages and tissues (Figure 2B,C). Although these interactions may not directly reflect immune or developmental activities, it suggests that the pleiotropic genes might participate in different processes by interacting with more molecular partners. The broader expression of pleiotropic genes across stages compared to non-pleiotropic genes suggests that one or both of the immune and developmental functions are required throughout ontogeny. Finally, among the most specifically-expressed immune genes (Figure 2B), pleiotropic genes were disproportionately expressed in embryos and pupae – key developmental stages – while the maximum expression of non-pleiotropic genes was more evenly distributed among post-embryonic life stages. This may reflect decoupling of immunological regulation across life stages, which could allow the different life stages to independently optimize immune responses over evolutionary time as they are exposed to different parasites and ecological conditions (Fellous and Lazzaro 2011; Critchlow, et al. 2019; Rolff, et al. 2019). In the future, it would be interesting to clarify the extent to which pleiotropic genes exhibit temporal segregation of developmental processes and immune roles in different life stages, as opposed to simultaneous participation in both functions in one or more stages.

Our results suggest a significant association between pleiotropy status and the rate of molecular evolution in immune system genes. Other studies that have considered the general relationship between signatures of molecular evolution and molecular pleiotropy have reached contrasting conclusions. In some cases, pleiotropy, as defined by connectivity in protein-protein or gene co-expression networks, is negatively correlated with molecular evolution rates (Alvarez-Ponce, et al. 2017; Masalia, et al. 2017) as we observe in our study. Meanwhile, others have detected very minimal or no correlation (Hahn, et al. 2004; Fraïsse, et al. 2018). The variance in these results could be attributed to differences in study organisms, different experimental contexts and the inherent differences in the various definitions of pleiotropy. For example, our definition of pleiotropy focused on two primary traits rather than considering the entire constellation of traits that might push estimates of pleiotropy in immune systems even higher. The two traits we chose, however, cover the extreme ends of evolutionary rate predictions, as development is thought to be one of the most conserved processes (Artieri, et al. 2009), while immunity is consistently identified as one of the most rapidly evolving systems across studied taxa (Obbard, et al. 2006; Areal, et al. 2011).

Our analyses of positive selection in pleiotropic and non-pleiotropic immune genes suggests that while the ω of positively selected sites is lower in pleiotropic immune genes, there are twice as many positively selected sites (1.5% vs 0.7% in PAML model M8 output) in pleiotropic immune genes relative to non-pleiotropic ones. Additionally, α values, which represent the proportion of substitutions drive by positive selection, were significantly higher in pleiotropic genes than in the other two categories. These results reflect key conclusions from a recent study demonstrating that virus-interacting proteins that participate in diverse cellular processes, which are otherwise more evolutionarily constrained, also showed higher rates of adaptation relative to those that are not known to interact with viruses (Enard, et al. 2016). We speculate that when mutations occur in pleiotropic proteins that have antagonistic effects on immunity or development, compensatory substitutions could arise to resolve this conflict. For example, a previous study suggested that the presence of a non-synonymous mutation greatly increases the chance of finding other substitutions nearby, possibly reflecting the correlated evolution of codons within a protein module (Callahan, et al. 2011). Because our analyses are not domain-specific, we cannot parse signatures of selection on regions within a pleiotropic gene that might provide specific immune or developmental functions or that could be closely associated with compensatory mutations. Although such analysis would require very specific knowledge of the effect of each mutation on immune and development phenotypes, future analyses could focus on a subset of genes with well-defined protein domain structures and protein-protein interaction data (e.g. Dredd and Jak; Figure 4, gold stars) to refine the functional and evolutionary significance of pleiotropic activity.

Across immune pathways, intracellular components are disproportionately pleiotropic compared to extracellular components (Figure 4). Interestingly, however, we observed that many pleiotropic intracellular signaling components associate with non-pleiotropic adapters or interact with proteins that exhibit higher rates of adaptation, which could provide a way to modify pleiotropic protein function in specific immunological contexts to relieve antagonism (Kinsler, et al. 2020). This analysis raises new questions for future investigation: how can a signaling pathway balance its role in multiple biological processes? What are the key players and their characteristics that affect how a pathway is used across several contexts or life stages?

Overall, our study serves as the first one to systematically quantify the degree of pleiotropy in a specific biological context and investigate correlations between pleiotropy and rates of molecular evolution in immune systems. These results lay the groundwork for future work to tease apart the mechanistic framework of these pleiotropic patterns to understand how genetic architecture shapes the mode and tempo of immune system evolution and their influence on immune phenotypes.

## Methods

### Immune and developmental gene list curation

We curated a comprehensive list of genes representing immunity by combining several resources, starting with manually curated list from previous immune studies (Lemaitre and Hoffmann 2007; Early, et al. 2017), which include most experimentally validated “canonical” immune genes. Separately, we appended Gene Ontology (GO)-annotated genes under the term “immune system process” (GO:0002376) to the list. We further sub-divided genes under this GO term into either “Immune Response” or “Immune Development” genes to differentiate between genes that play direct roles in mounting an immune response and genes contributing to the development and maturation of the immune system. Finally, we added to our list a core set of immune genes from (Troha, et al. 2018), which comprises 252 genes that show differential expression across infection with ten different bacterial species of variable virulence.

For each immune gene, we also assigned an immune gene class – recognition, signaling, or effector - based on the gene’s known function in the immune system. If a gene has not been assigned a class in previous studies, we manually assign it a class based on the gene description from FlyBase. For a detailed description of each gene class definition, see Supplemental Protocol.

Separately, we created a list of GO-annotated developmental genes by querying the term “Developmental Process” (GO:0032502), while separately annotating genes belonging to the child term “embryonic morphogenesis” (GO:0048698). All GO annotation queries were conducted through FlyBase (Thurmond, et al. 2019). A full list of genes in each group is included in Supplemental Table 1 and visualization of the degree of overlap between different resources is in Supplemental Figure 1.

### Pleiotropy categorization

Pleiotropy refers to the phenomenon where a single gene influences multiple traits. However, the definition of “trait” can be ambiguous across different biological contexts, and thus pleiotropy can manifest at different levels and be detected by various methods (Paaby and Rockman 2013; Tyler, et al. 2016). At the molecular level, pleiotropy can refer to the multiple biochemical roles that a gene can have and is frequently measured as the number of physical interacting partners (Hahn, et al. 2004). At the developmental or phenotypic level, pleiotropy can involve genes affecting distinct phenotypes or biological processes, as measured by the number of stage or tissues in which such genes are expressed (Artieri, et al. 2009). Lastly, under an evolutionary perspective, pleiotropy can refer to the separate components of fitness that a gene might modulate, a well-known example being the antagonistic pleiotropy model for the evolution of aging (Williams 1957). Though many interpretations of pleiotropy exist, in this study, we are specifically concerned about pleiotropic genes at the phenotypic level. In particular, we focused on genes annotated to play roles in both immune and developmental processes. As such, if a gene is annotated as functioning in both immunity and development from the lists curated from the method described above, it was considered pleiotropic. A full list of pleiotropic genes is included in Supplemental Table 1.

For comparison purposes, we also calculated molecular metrics of pleiotropy for each gene in the genome regardless of annotated function in immunity or development. These measurements include expression stage specificity (described below), number of associated Biological Processes GO terms, number of associated Molecular Functions GO terms, number of protein-protein interactions, and number of gene-gene interactions. All raw data files were obtained through the FlyBase ftp server, and the latest version of each file was downloaded (March 2020, Supplemental Protocol).

### Categorization of stage and tissue specificity

Genes with functions limited to specific tissues or life stages (and particularly later life stages) may have less pervasive effects on organismal fitness (Cutter and Ward 2005; Artieri, et al. 2009), possibly buffering evolutionary constraint from pleiotropy. To calculate expression specificity, we applied the following equation (Yanai, et al. 2005) to expression level data of all *D. melanogaster* genes in all stages (embryo, larva, pupa, adult) and tissues (Supplemental Methods):

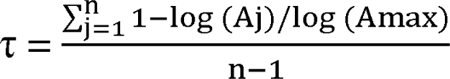

In this equation, n is the number of of stages or tissues. A_j_ is the expression level at stage/tissue j, A_max_ the maximum expression level of of stages/tissues. Lower tau (τ) values signify specific expression in a certain stage/tissue, while a higher one indicates broad expression across all stages/tissues (Fraïsse, et al. 2018). Tau values for all of the genes used in the analysis are provided in Supplemental Table 6.

### Pathway annotation

We used the PANTHER database to annotate our gene lists to pathway, if available. In short, all genes are compiled into a list of IDs, which is then used as a query in PANTHER (http://pantherdb.org/). We then downloaded the annotations and computed the total number of unique pathways associated with each gene group (pleiotropic vs. non-pleiotropic).

### Compiling sequences for PAML analyses

Genes included in our analyses were chosen using the Table 1 row 2 inclusion criteria for non-pleiotropic immune (454 genes), pleiotropic (299 genes), and non-pleiotropic developmental (3047 genes) lists. We used the FlyBase gene IDs to download coding sequences (CDSs) using the FlyBase Sequence Downloader tool (FB2021_05, released October 15, 2021) for *D. melanogaster* (Thurmond, et al. 2019). We then obtained a list of orthologs from FlyBase for all 12 sequenced *Drosophila* species. Using custom scripts (https://github.com/alissawilliams/pleiotropy_Drosophila/tree/main/scripts), we parsed out FlyBase sequence IDs for 11 other *Drosophila* species (Supplemental Table 5) for the genes of interest using the *D. melanogaster* IDs. We used the Sequence Downloader tool from an archived version of FlyBase (FB2017_05, released October 25, 2017) to download CDSs for each gene of interest for each of the other 11 species.

We used another set of custom scripts to compile one sequence file for each gene of interest within each pleiotropy category. These scripts added one CDS per species to each file; in cases where more than one CDS was obtained for a single gene ID, the first CDS in the file of downloaded sequences was used. In cases of paralogy (i.e. where one species had multiple gene identifiers within a single orthogroup), the species with gene duplicates were excluded from the sequence file. After this step, 400, 294, and 2549 sequence files contained at least two sequences for the non-pleiotropic immune, pleiotropic, and non-pleiotropic developmental groups, respectively.

Next, sequence files containing at least two sequences were aligned in codon space with the *einsi* option in MAFFT v7.310 (Katoh and Standley 2013) using a custom script (https://github.com/dbsloan/perl_modules). Successful alignment occurred for 356 non-pleiotropic immune genes, 231 pleiotropic genes, and 2067 non-pleiotropic developmental genes. These alignment files were trimmed in codon space using Gblocks v0.91b (Castresana 2000) with parameters *-t = c* and *-b5 = h*. These trimmed files were used in downstream PAML analyses.

### Calculating gene-wide d_N_/d_S_ values using PAML

The trimmed sequence files were individually run through codeml site model M0 in PAML v4.9j (Yang 2007) to obtain *d_N_/d_S_* values for each gene. The codeml command was run using *seqtype = 1*, *CodonFreq = 2*, *model = 0*, *NSsites = 0*, and *cleandata = 0*. Constraint trees for each gene were built by starting with the known species tree for the 12 *Drosophila* species on FlyBase and eliminating any species not present in the particular sequence file. The site model M0 runs were successful for 348 of the 356 non-pleiotropic immune genes, 227 of the 231 pleiotropic genes, and 2037 of the 2067 non-pleiotropic developmental genes. *d_N_/d_S_* values across the three gene categories were compared using a Kruskal-Wallis test followed by post-hoc Dunn tests in R (R_Core_Team 2012).

### Detection of positive selection using PAML site models

To detect positive selection in genes of the three categories, we used codeml site models M7 and M8 in PAML. The trimmed files for each category were concatenated into single alignments and run through codeml with parameters *seqtype = 1*, *CodonFreq = 2*, *model = 0*, *NSsites = 7 8*, and *cleandata = 0*. A constraint tree for the 12 *Drosophila* species was built based on the phylogeny provided on FlyBase (Thurmond, et al. 2019). Within each class of genes, models M7 and M8 were compared using likelihood ratio tests (df = 2). Site model M0 (*model = 0, NSsites = 0)* was also run for each of the three concatenated gene sets using the same parameters as described in the previous section. Comparisons of proportions of sites under positive selection were calculated in two ways. First, the proportion identified by model M8 was multiplied by the total number of sites in each concatenated alignment and rounded to the nearest whole number for use in a Chi-squared test in R. Pairwise Chi-squared tests were also conducted and the p-value used for detecting significance was determined using a Bonferroni correction for multiple testing. Second, we extracted *d_N_/d_S_* values from the Bayes Empirical Bayes analysis output (also from model M8) for all sites with a reported probability of *d_N_/d_S_ >* 1 of 0.95 or above. The numbers of sites under positive selection using this criterion were also used in Chi-squared tests, along with the total number of sites in each alignment. Again, pairwise Chi-squared tests in R were evaluated after Bonferroni correction for multiple testing.

In addition to the concatenated sequences, we ran codeml site models M7 and M8 on individual pleiotropic and non-pleiotropic immune genes from the three KEGG-annotated immune signaling pathways (Figure 4).

### Calculation of α and ω_a_ using MultiDFE

To calculate the proportion of substitutions driven by positive selection (α) and the rate of adaptive substitutions (ω_a_), we used PopFly data from the Raleigh (RAL) and Zambia (ZI) populations (Hervas, et al. 2017) in the iMKT package in R (Murga-Moreno, et al. 2019) as input to the software package MultiDFE (https://github.com/kousathanas/MultiDFE). The MultiDFE input was in the form of site frequency spectra (SFS). The PopFly data was obtained from the file dsimDmelSites.tab provided by Jesús Murga-Moreno (Murga-Moreno, et al. 2019). Of the 356 non-pleiotropic immune genes, 231 pleiotropic genes, and 2067 non-pleiotropic developmental genes included in the concatenated alignments, the dsimDmelSites.tab contained 317, 207, and 1757, respectively, for the RAL population and 350, 226, and 1959, respectively, for the ZI population. We modified code in the iMKT Jupyter notebook (https://nbviewer.org/github/jmurga/iMKTData/blob/master/notebooks/dmelProteins.ipynb, accessed 1 June 2022) to obtain raw counts of variants for each gene in each population. We then used bootstrapping to create 100 samples for each gene class in each population by summing variant counts as well as pi, p0, di, d0, mi, and m0 from the iMKT PopFlyData table (Murga-Moreno, et al. 2019). We calculated the 0^th^ column of each SFS (i.e. the number of sites with no observed variants) using the equations mi – pi and m0 – p0 for nonsynonymous and synonymous sites, respectively. Scripts used for this process are provided at https://github.com/alissawilliams/pleiotropy_Drosophila/tree/main/scripts.

We ran MultiDFE with the recommended parameters *-conpop 0, -sfsfold, 1 -selmode 4, -nspikes 0,* and *-ranrep 1* (Kousathanas and Keightley 2013) for each bootstrapped SFS file (https://github.com/kousathanas/MultiDFE, downloaded 14 April 2022). We then extracted the average fixation probability (fix_prob) for each bootstrap replicate for each population from its respective .sfs.MAXL.out output file and calculated α and ω_a_ by plugging fix_prob from MultiDFE and the summed di and d0 from PopFlyData into equations 10 and 11 from (Kousathanas and Keightley 2013). Values of di and d0 were corrected using the Jukes-Cantor correction function provided on the MultiDFE GitHub page (https://github.com/kousathanas/MultiDFE, accessed 14 April 2022). Distributions of α and ω_a_ values were compared for the RAL and ZI populations separately using a Kruskal-Wallis test followed by post-hoc Dunn tests in R (R_Core_Team 2012) in cases where the Kruskal-Wallis test produced a significant result.

## Statistical Analysis

All statistical analyses were conducted in R (4.1.0). We used Shapiro tests to assess distribution normality in datasets. For comparison between multiple groups, we conducted Kruskal-Wallis tests followed by pairwise Dunn tests with Benjamini-Hochberg correction in cases where there was a significant difference between groups.

## Acknowledgments

We thank Nora Schulz, Stephanie Birnbaum, and other members of the Tate lab for comments and discussion. We also thank Seth Bordenstein and two anonymous reviewers for their helpful comments on an earlier version of this manuscript. Additionally, we thank Athanasios Kousathanas for his help with compilation and use of the MultiDFE software and Jesús Murga-Moreno for his help with the iMKT package. This work was supported by the National Institute of General Medical Sciences at the National Institutes of Health (grant number R35GM138007 to A.T.T.).

## Data Availability

Gene classifications, scripts, untrimmed and trimmed alignments, PAML output, and MultiDFE input and output are provided at https://github.com/alissawilliams/pleiotropy_Drosophila. Additional data are provided in the Supplemental Tables and Figures.

## List of Supplemental Tables and Figures

**Supplemental Table 1:** Categorization of each gene included in analysis according to different definitions of developmental pleiotropy as described in Table 1

**Supplemental Table 2:** Genes manually annoted for function derived from FlyBase (see Methods; Comparison of pleiotropic and non-pleiotropic immune gene characteristics)

**Supplemental Table 3:** Assignment of pleiotropic and non-pleitropic genes to PANTHER pathways ntal Table 3:

**Supplemental Table 4:** Corresponding number of paralogs among Drosophila species for each Dmel gene

**Supplemental Table 5:** The number and percentage of specific pleiotropic and non-pleiotropic genes that showed maximum expression in each stage

**Supplemental Table 6:** Full dataset used in statistical analysis of tau results

**Supplemental Figure 1:** Venn Diagram representing the overlap between sources used to curate the immune gene list.

**Supplemental Figure 2:** Venn Diagram representing the overlap between sources used to curate the developmental gene list.

**Supplemental Figure 3:** Number of pleiotropic and non-pleiotropic genes in each immune gene list.

**Supplemental Figure 4:** Number or Biological Processes and Molecular Function GO terms associated with genes belonging to each pleiotropy group.

**Supplemental Figure 5:** Values of ω_a_ for each of the three gene categories in both the Raleigh (RAL) and Zambia (ZI) populations of *Drosophila melanogaster*.

