## Supplemental Figures for "The effect of developmental pleiotropy on the evolution of insect immune genes"

T.M. Ngo, A.M. Williams, and A.T. Tate

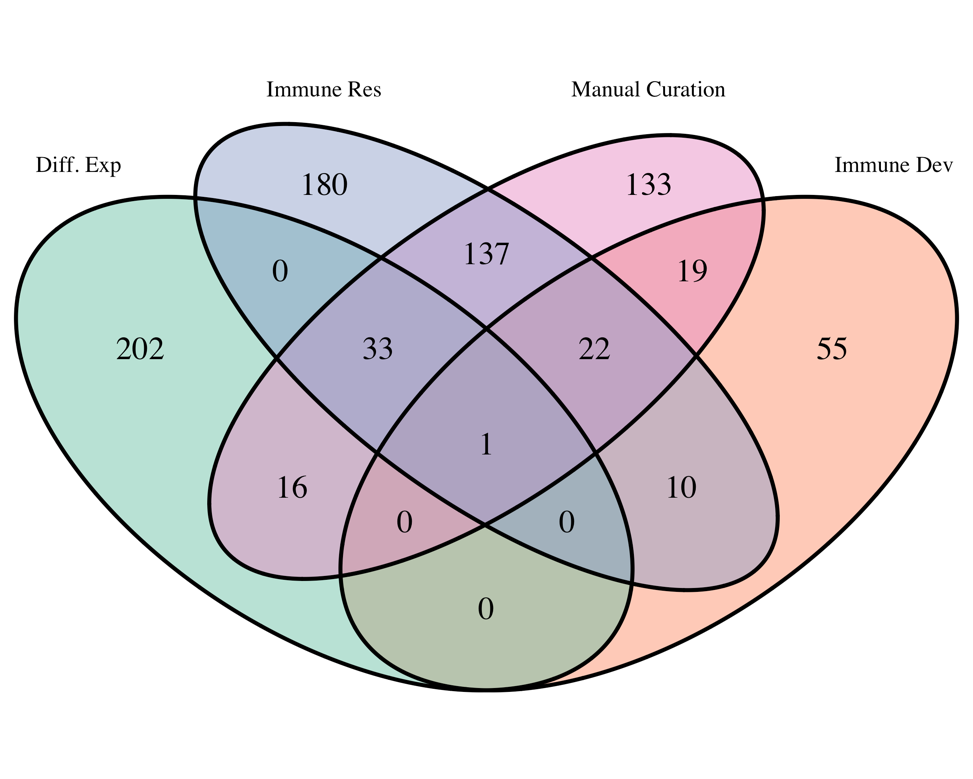

**Supplemental Figure 1**: Venn Diagram representing the overlap between sources used to curate the immune gene list. Four distinct lists are combined to obtain a list of 808 immune genes:

1. “Diff Exp” group refers to the core set of differentially expressed gene identified in Troha et al 2018.
2. “Immune Res” group refers to genes annotated under the following GO Terms that represent the core actions of an immune response:
   a. GO:0002252 Immune Effector Process
   b. GO:0002253 Activation Of Immune Response
   c. GO:0006955 Immune Response
   d. GO:0019882 Antigen Processing
   e. GO:0045321 Leukocyte Activation
   f. GO:0035172 Hemocyte Proliferation

Each GO term is queried in FlyBase and all six lists are concatenated, and duplicates are removed.

3. “Manual Curation” group refers to a manually curated immune gene in Early et al 2018, which also cites previous experimental studies on *Dmel* immunity.
4. “Immune Dev” group refers to genes annotated under the following GO Terms that represent genes having a role in the *development* of the immune system instead of direct participation in the immune response:
   a. GO:0002520 Immune System Development
   b. GO:0042386 Hemocyte Differentiation

Each GO term is queried in FlyBase and all six lists are concatenated, and duplicates are removed. The detailed description of each GO Term can be viewed in FlyBase.

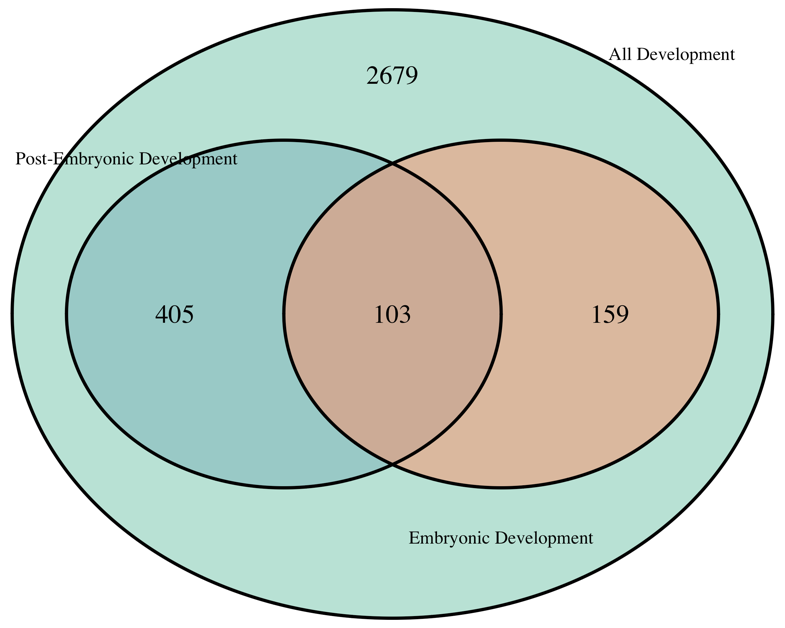

**Supplemental Figure 2:** Venn Diagram representing the overlap between sources used to curate the developmental gene list. Similar to the method that we used to curate the immune gene list described above, three separate GO Terms are queried through FlyBase and the list of developmental gene is annotated in accordance to their GO Term annotation:

1. GO:0032502 Developmental Process
2. GO:0048598 Embryonic Morphogenesis
3. GO:0009886 Post-embryonic Development

Throughout the study, “developmental gene” refers to all 3346 genes, unless otherwise noted. The detailed description of each GO Term can be viewed in FlyBase.

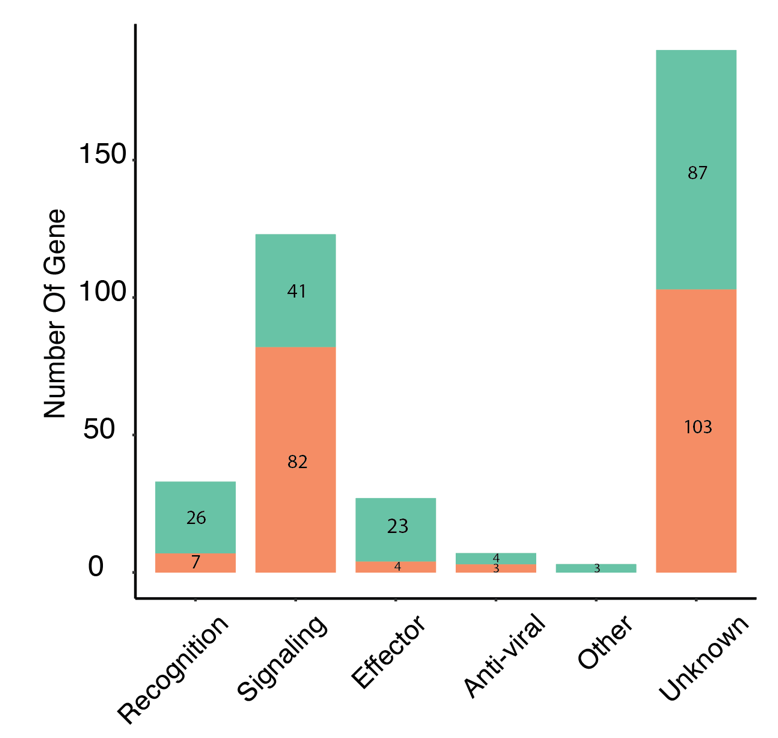

**Supplemental Figure 3:** Number of pleiotropic (orange) and non-pleiotropic (green) genes in each immune gene list.

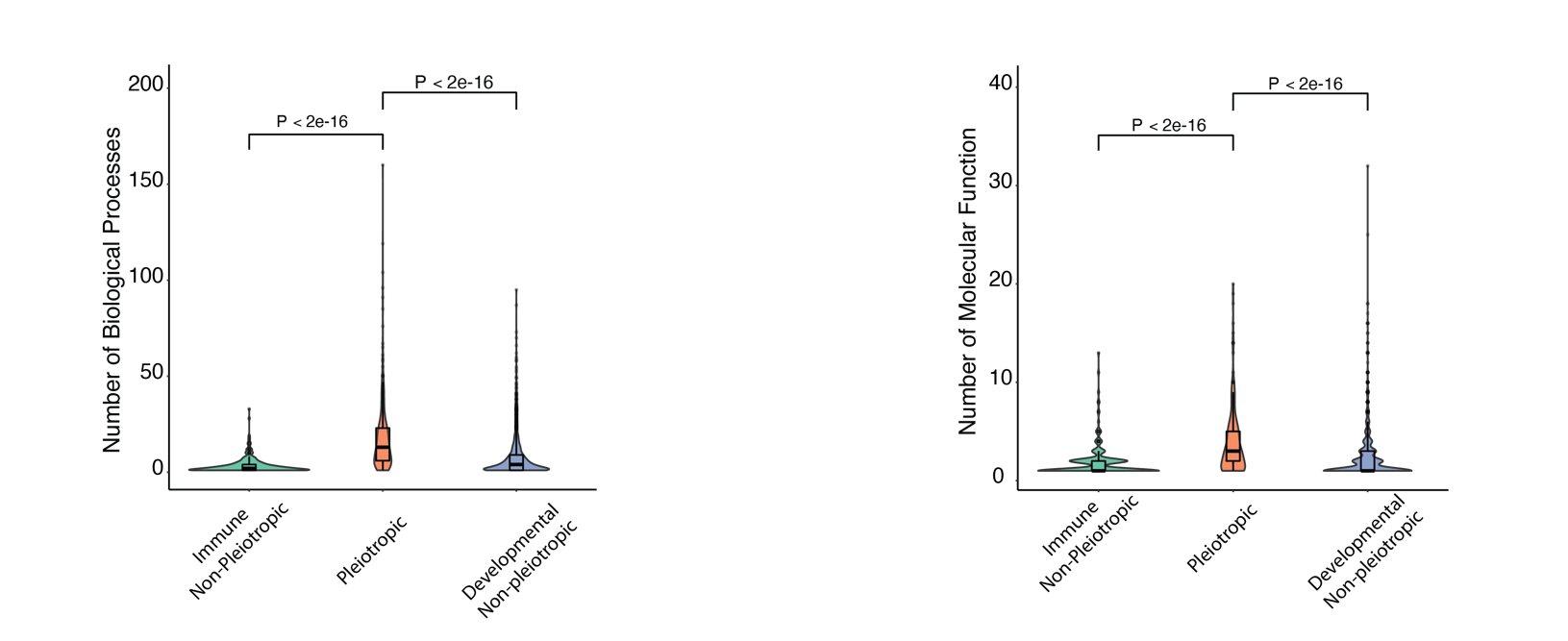

**Supplemental Figure 4:** Number or Biological Processes and Molecular Function GO terms associated with genes belonging to each pleiotropy group.

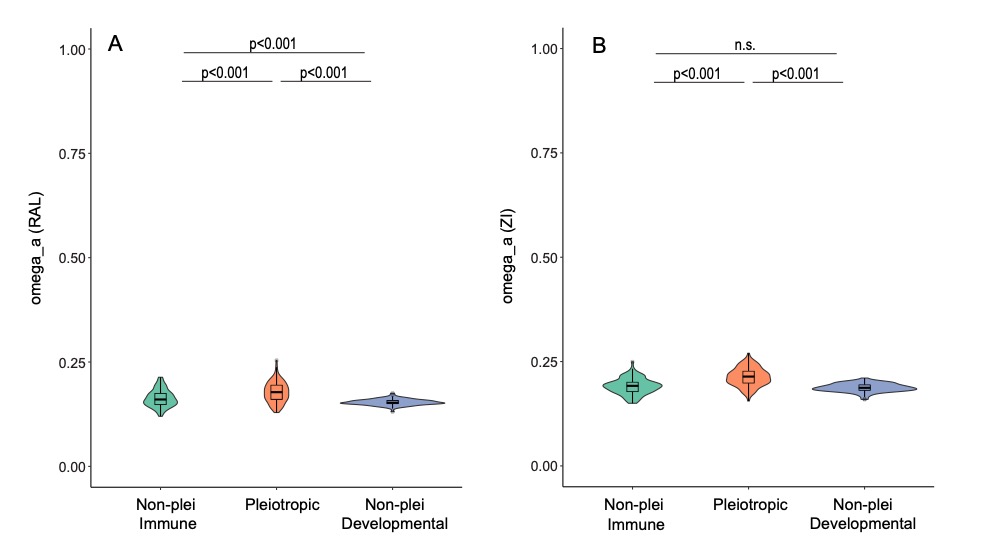

**Supplemental Figure 5:** Values of *ω_a_* for each of the three gene categories in both the Raleigh (RAL) and Zambia (ZI) populations of *Drosophila melanogaster*. *ω_a_* values were calculated using MultiDFE on 100 bootstrap replicates of summed site frequency spectra (SFS) for each gene category. Distributions were compared using a Kruskal-Wallis test followed by post-hoc Dunn tests in R. Sample sizes are identical to those in Fig. 3C and D.
